# SATurn: A modular bioinformatics framework for the design of robust maintainable web-based and standalone applications

**DOI:** 10.1101/313619

**Authors:** David R. Damerell, Claire Strain-Damerell, Sefa Garsot, Stephen Joyce, Paul Barrett, Brian D. Marsden

## Abstract

**Summary:** SATurn is a modular, open-source, bioinformatics platform designed to specifically address the problems of maintenance and longevity commonly associated with the development of simple tools funded by academic research grants. Applications developed in SATurn can be deployed as web-based tools, standalone applications, or hybrid tools which have the benefits of both. Within the Structural Genomics Consortium (SGC) we have utilized SATurn to create a bioinformatics portal which routinely supports a diverse group of scientists including those interested in structural biology, cloning, glycobiology & chemicalbiology.

**Supplementary information:**

## 1 Introduction

The majority of bioinformatics tools and databases developed in academia are funded by fixed-term responsive mode grant applications and as such are commissioned as research projects (Nature Methods (Editorial) 2016). Ensuring sustainability for tools and databases which are outside of their innovative phase and thus no longer eligible for the majority of research grants is a stumbling block for many such resources. For example, KEGG (Kyoto Encyclopedia of Genes and Genomes) (Kanehisa, Sato et al. 2016) is a particularly well-known resource which suffered from this funding phenomenon (Kanehisa 2014).

Poorly maintained computational tools and databases present two challenges to scientists. Firstly, useful tools and databases may simply become unavailable. Secondly, those tools and databases which are not maintained may provide at best incomplete and at worst misleading outputs if they do not incorporate recent data-sets.

To tackle the first problem many funding bodies and publishers have instigated policies which demand that tools and databases are made available for at least a given fixed-term (typically 2 years) after publication and that source code is made freely available. In practice it can be challenging for researchers to guarantee that a resource will be available for such a fixed-term due to both a lack of ongoing funding and the likelihood that the original developers of the resource will have moved elsewhere.

Popular bioinformatics tools often share the characteristics of being simple to use and bestowing increased efficiencies or insights to the end user. The ExPASy web-site (Gasteiger, Gattiker et al. 2003) is an example of a well-known set of accessible tools used by biochemists and molecular biologists.

Bioinformaticians have been aware of the issues surrounding software sustainability for a great many years and have strived to create technologies that can support the development of maintainable and reusable software. Repositories of reusable libraries and plugins which others can use to build applications are a commonly employed to address, to a degree, the issue of maintainability. BioJS is one such repository for JavaScript Bioinformatics libraries (Gomez, Garcia et al. 2013). Although extremely useful, such repositories do not directly address the problem of how to ensure web-applications, built upon libraries, remain available to the community when they are no longer available for download from the authors’ own web-site. Contemporary technologies (such as Docker) make it possible for web-applications to be provided as pre-configured, self-contained, containers or images. The Galaxy platform (Afgan, Baker et al. 2016) has perhaps made the greatest strides in improving the maintainability and sustainability of Bioinformatics programs by encouraging developers to contribute programs and workflows to a common platform. Galaxy also provides Docker images for installation

Technologies built around the Jupyter Notebook environment make it easy for developers of all abilities to create simple scripts which they can easily share with others. However users hoping to take advantage of shared Jupyter notebooks must install software dependencies themselves. These software dependencies drastically increase the long-term sustainability burden, and may render the notebook itself useless after development effort has ended.

At the SGC we wanted to create a new Bioinformatics portal for our scientists whilst attempting to avoid the issues so far outlined as far as possible. Whilst recognizing that many existing technologies discussed do address aspects of maintainability and sustainability none of them focus on providing users with a simple user interface and application which can be deployed locally or remotely in a sustainable fashion. Described herein is the SATurn platform and the Bioinformatics portal developed within it.

## 2 SATurn Platform

SATurn is a bioinformatics platform which overcomes the issues discussed above regarding software sustainability, maintainability, and extendibility. SATurn provides a common user interface (UI) and simple code environment which can be used to design diverse bioinformatics tool plugins. Multiple plugins can be served from the same SATurn instance allowing developers to create bioinformatics portals based upon local workflows rapidly and with minimal effort. SATurn portals can be deployed as standalone desktop applications or remotely hosted web-portals. The SATurn framework has been used to develop a bioinformatics portal for SGC scientists, now in use for over three years, which simplifies many common bioinformatics tasks associated with medium/high throughput cloning. Tools are included for viewing protein & DNA sequences, sequencing files, compound structures and glycans. Phylogenetic trees, sequence summary figures, and multiple sequence alignments can also be generated. See section 4 of the supplementary materials for an example of a program included. This bioinformatics portal can be downloaded for Windows, OS/X, and Docker from: https://github.com/ddamerell53/SATurn/releases (see section 2 of the supplementary materials). Developers can get started with SATurn by following the instructions on the SATurn wiki: https://ddamerell53.github.io/SATurn/.

### 2.1 Architecture

SATurn portals consist of a web-based UI (see section 3 of the supplementary materials) and a server-side component which runs inside a NodeJS web-server (https://nodejs.org). The architecture of the SATurn platform has been specifically developed with a number of key features to improve the sustainability, maintainability, and extendibility of plugins and portals created using the platform.

To overcome the problem of hosted web-applications becoming unavailable over time SATurn portals can be deployed as both standalone applications and web-portals. The SATurn platform has three deployment modes which are shown in Table 1. This ensures that any portal is available for use on a desktop computer even if it is no longer provided as a web-based application.

**Table 1:** SATurn deployment modes

| Mode | Web-server | Client Access | Access Files |
| --- | --- | --- | --- |
| **Fully Hosted** | Remote | Web-browser | No |
| **Standalone** | Local | Application | Yes |
| **Hybrid** | Remote | Application | Yes |

Many of the most useful academic bioinformatics tools are created by wet-laboratory scientists for their own use and as such have little, if any, formal training in software development. We have therefore designed SATurn specifically with such developers in mind and have made it extremely simple to add their tools as plugins to the platform. The SATurn UI includes a built-in code editor which can be used to design new plugins, extend existing ones and write simple scripts to analyze data. SATurn includes a simple way of enabling two-way communication between SATurn and tools which have already been created in languages like Python and PERL (see section 5 of the supplementary materials for an example).

## 3 Conclusion

Being able to invest an appropriate degree of time and resource into building useful bioinformatics tools that can stand the test of time is not a luxury that is always available to all. This results in many initially useful tools becoming prematurely unavailable. Recognizing these constraints, we have created a new extendable platform, SATurn, within which bioinformatics tools can be created for use as standalone desktop applications as well as web-sites. SATurn represents a fundamental tool in many aspects of the work performed at the SGC and is also a critical component in a range of training events. We continue to add new functionality to the platform with a view to providing useful functionality beyond basic bioinformatics capabilities. Whilst the SGC is committed to maintaining the SATurn platform, we have deliberately developed it with the intention of providing an environment which makes it as likely as possible that the software embedded within the platform will continue to work for years to come.

## Acknowledgements

The SGC is a registered charity (number 1097737) that receives funds from AbbVie, Bayer Pharma AG, Boehringer Ingelheim, Canada Foundation for Innovation, Eshelman Institute for Innovation, Genome Canada through Ontario Genomics Institute [OGI-055], Innovative Medicines Initiative (EU/EFPIA) [ULTRA-DD grant no. 115766], Janssen, Merck KGaA, Darmstadt, Germany, MSD, Novartis Pharma AG, Ontario Ministry of Research, Innovation and Science (MRIS), Pfizer, São Paulo Research Foundation-FAPESP, Takeda, and Wellcome [106169/ZZ14/Z]. We would also like to acknowledge the Oxford Protein Production Facility staff who provided many fruitful discussions and contributed a small amount of code to the framework.

*Conflict of Interest:* none declared.

